# Atomistic simulations of RNA duplex thermal denaturation: sequence- and forcefield-dependence

**DOI:** 10.1101/2023.09.29.560124

**Authors:** Aimeric Dabin, Guillaume Stirnemann

**Affiliations:** CNRS Laboratoire de Biochimie Théorique, Institut de Biologie Physico-Chimique, Université de Paris Cité, 13 rue Pierre et Marie Curie, 75005, Paris, France; PASTEUR, Département de chimie, École normale supérieure, PSL University, Sorbonne Université, CNRS, 75005 Paris, France

## Abstract

Double-stranded RNA is the end-product of template-based replication, and is also the functional state of some biological RNAs. Similarly to proteins and DNA, they can be denatured by temperature, with important physiological and technological implications. Here, we use an *in silico* strategy to probe the thermal denaturation of RNA duplexes. Following previous results that were obtained on a few different duplexes, and which nuanced the canonical 2-state picture of nucleic acid denaturation, we here specifically address three different aspects that improve our description of the temperature-induced dsRNA separation. First, we investigate the effect of the spatial distribution of weak and strong base-pairs among the duplex sequence. We show that the deviations from the two-state dehybridization mechanism are more pronounced when a strong core is flanked with weak extremities, while duplexes with a weak core but strong extremities exhibit a two-state behavior, which can be explained by the key role played by base fraying. This was later verified by generating artificial hairpin or circular states containing one or two locked duplex extremities, which results in an important reinforcement of the entire HB structure of the duplex and higher melting temperatures. Finally, we demonstrate that our results are little sensitive to the employed combination of RNA and water forcefields. The trends in thermal stability among the different sequences as well as the observed unfolding mechanisms (and the deviations from a two-state scenario) remain the same regardless of the employed atomistic models. However, our study points to possible limitations of recent reparametrizations of the Amber RNA forcefield, which sometimes results in duplexes that readily denature under ambient conditions, in contradiction with available experimental results.

## Introduction

Nucleic acids have the ability to self-interact through hybridization, where one strand portion can interact with another strand portion through intermolecular hydrogen-bonds (HB). Desoxyribose nucleic acids (DNA) thereby form double-stranded (ds) duplexes, their native configurations in most of the biological contexts *in vivo*, while single-stranded (ss) RNA typically fold into three-dimensional structures that contain, but are not limited to, hybridized regions. Although rare in nature, dsRNA are involved in a variety of biological processes. In eukaryotes, most of them are related to the regulation of gene expression.^1–4^ In RNA-based viruses, dsRNA stores the virus’ genetic information. In the context of the RNA World, a strong and popular hypothesis to explain the origins of life, the first forms of life would have been formed by self-replicative oligonucleotide networks, with transient dsRNA products following template-based replication. ^5^

In cells, nucleic acid duplexes are typically separated by enzymes, for example during transcription for dsDNA. However, similarly to proteins that denature upon temperature-increase, nucleic acid strands that are hybridized can separate at high temperature.^6–8^ *In vivo*, some regulatory systems are based on the temperature-induced local deformation of base-paired motifs in so-called RNA thermoswitches used to regulate translation. ^9–12^ For technological applications within the framework of RNA origami and nanotechnology,^13,14^ temperature could also be an important factor governing the device assembly and disassembly.

Protein and nucleic acid melting curves (in the experiments as well as in the simulations) are most often interpreted within the framework of a two-state model, one state corresponding to the native, folded (or hybridized) conformation and the other one to an unfolded/separated version of the biomolecule. ^6–8,15–19^ From a free-energy landscape perspective, this would thus correspond to two free-energy basins whose coordinates in the configuration space are fixed but whose respective weights change with temperature. For RNA duplexes, which are the main focus of the current contribution, such a two-state model is also used to calibrate nearest neighbors models^7,15,20,21^ that enable the prediction of a given duplex melting temperature depending on its sequence, often with very good accuracy.

However, reports have been made, both from experimental studies^22–24^ as well as from theoretical considerations (some based on nearestneighbor models themselves), ^15,20,21^ of significant deviations from a two-state model for RNA duplexes (de)hybridization, with base fraying at the duplex extremities^25–31^ and the presence of stable intermediates.^32,33^ Similar observations were made, and were actually much better characterized, for DNA duplexes.^34–37^ In particular, molecular dynamics (MD) simulations were instrumental in gaining a molecular picture of the associated molecular mechanisms, mostly using coarse-grained representations. ^35,38–41^ Because of the limitations in the phase space exploration offered by unbiased, all-atom MD simulations in explicit solvent,^42,43^ enhanced sampling strategies along the temperature ladder or along well-identified collective variables (CVs) offer important mechanistic insight.^44,45^

In a very recent work, we deployed such an enhanced sampling strategy with an all-atom biomolecular representation in explicit solvent to unravel the molecular details of RNA thermal dehybridization. ^46^ While we had previously shown that this approach was successful to study the thermal denaturation of proteins,^18,19,47–50^ we demonstrated that, for a selection of experimentally-available short duplex structures with either A-U or G-C base pairs, as well as for artificial structures with random sequences, melting curves and melting temperatures in very good agreement with available experimental data and nearest-neighbor models could be obtained.^46^ We then showed that the unfolding mechanism of the duplexes was more subtle than that of a two-state picture, with significant base-fraying at the extremities, that progressively propagates from the duplex extremities to its core as temperature increases.^46^ This gave rise, for all the investigated sequences, to intermediates states around melting that were structurally very distinct from the fully-formed duplexes. As a consequence, temperature results in a progressive shift in the duplex structure, rather than a progressive depletion of a stable duplex state and population of a separated, dehybridized state.^46^

Our initial study was limited to duplexes containing either weak A-U base pairs, or strong G-C base pairs, or, complementary duplexes with random sequences, and it was based on one given RNA/water forcefield combination for most investigated structures. In this contribution, we extend and we improve our investigation of the temperature-induced dsRNA separation. First, we investigate the effect of the spatial distribution of weak and strong base-pairs among the duplex sequence, by generating artificial structures that contain a fix 50:50 ratio of A-U and G-C base pairs, but with distinct patterns. We show that the deviations from the two-state dehybridization mechanism are more pronounced when a strong core is flanked with weak extremities, while duplexes with a weak core but strong extremities exhibit a two-state behavior. However, a scenario whereby the core would melt before the extremities is never observed. Second, from experimentally available structures containing either A-U or G-C, we generate artificial hairpin or circular states by constraining one or two duplex extremities. By suppressing the weak points of the duplex denaturation, we observe significant shifts in the duplex thermal stability, which results from an important reinforcement of the entire HB structure of the duplex.

A third and important aspect that we investigate is whether these results depend on the employed forcefields for RNA and water. We show that while these affect the exact values of the simulated melting temperatures, the trends in thermal stability among the different sequences as well as the observed unfolding mechanisms (and the deviations from a two-state scenario) remain the same regardless of the employed atomistic models. However, our study points to possible limitations of a recent and widely used state-of-the-art reparametrization of the Amber99 forcefield, which sometimes results in duplexes that readily denature under ambient conditions, in contradiction with available experimental results.

## Methods

### Systems and molecular models

In order to investigate the effect of strand sequences on the melting properties, we compare artificial dodecamer duplex systems (“art” prefix). Initial structures were generated using the X3DNA web interface. ^51^ We first chose two sequences containing either A-U or G-C base-pairs only: 5’-UAUAUAUAUAUA-3’ (art-(UA)_6_) and 5’-CGCGCGCGCGCG-3’ (art-(CG)_6_); then 3 sequences (and their complementary strands) with an equal amount of A-U and G-C base pairs but different motifs: 5’-GGGAAAAAAGGG-3’ (art-G_3_A_6_G_3_), 5’-AAAGGGGGGAAA-3’ (art-A_3_G_6_A_3_), and 5’-AGAGAGAGAGAG-3’ (art-(AG)_6_). Finally, we generated two random sequences and their complementary strands: 5’-UAGGAGGCUAGA-3’ (art-RND_1_) and 5’-GAACUGUAUGAU-3’ (art-RND_2_).

Missing hydrogens were added using the Pdb2gmx module from Gromacs package^52^ based on the residue templates of the employed forcefield (see below). Each double-stranded RNA was solvated with about 6,700 water molecules using the TIP3P water model geometry such that to fit in a box of 60x60x60 Å^3^ and Na^+^ cations were added to neutralize each system. These simulations in neutral conditions mean that the comparison with nearest-neighbor model predictions (at 1M NaCl) are expected to differ by 5-10 K, which corresponds to the shift in melting temperatures typically seen in the experiments when comparing low and high salt concentrations. ^53^ There are many other possibles sources of discrepancies and errors that could in particular arise from the forcefield that such a small difference is not relevant here; note however that we have demonstrated in our previous study that our approach is able to capture the dependence of the duplex stability upon ion concentration.^46^

The bulk of our simulations was propagated using the parameters of the ff99 forcefield with bsc0 and *χ*_OL3_ corrections^54^ for the RNA; for sodium, the Joung-Cheatham parameters; ^55^ and for water, the TIP3P forcefield.^56^ This combination is widely used and typically offers a robust description of the RNA structural and dynamical properties.^25^ In order to investigate the potential sensitivity of our results towards a change in forcefield, we tested the following alternatives. First, we varied the water model only, using the SPC/E^57^ or the OPC^57^ water models instead of TIP3P.^56^ Another set of simulations were performed using recent corrections of our original RNA forcefield (*ϵ ζ*_OL1_ alone or together with *β*_OL1_ ^58,59^), in TIP3P water, or the very recent gHBfix21 potentials^60^ with the original *χ*_OL3_ forcefield^54^ including vdW phosphate corrections,^61,62^ in OPC.^63^

To investigate the impact of base fraying on the unfolding, we artificially constrained the hydrogen bonds of one extremity (E1) or two (E2). We applied upper wall harmonic potentials using the PLUMED version 2.8,^64^ with a force constant of 500 kJ.mol^−1^ and a center which depends on the considered HB (Table 1), to allow for some fluctuations compared to their crystal structure values, but preventing them from total separation.

**Table 1:** Hydrogen-bond (HB) distances threshold used for the contraints of a double-strand RNA extremity.

|  |  |  |
| --- | --- | --- |
| Uridine-Adenine | O4-H61 | 2.5 Å |
|  | H3-N1 | 2.4 Å |
| Cytosine-Guanine | H42-O6 | 2.4 Å |
|  | N3-H1 | 2.4 Å |
|  | O2-H22 | 2.3 Å |

A timestep of 2 fs was employed, with a cut-off of 1.2 Å for Lennard-Jones interactions, and a short-range electrostatic cut-off of 1.2 Å. Long-range electrostatic interactions were treated using the Ewald summation^65^ on a 1.2 Å grid. All simulations were performed with the Gromacs (2019.2 or 2022.4 versions).^52^

### Hamiltonian replica exchange with solute scaling (REST2)

Each system was first minimized using a steepest decent algorithm, and then equilibrated using a Berendsen thermostat (target: 300 K) and barostat (target:1 atm) for 2 ns (during the first 1 ns, the biomolecule positions were restrained using force constants of 1000 kJ.mol^−1^.nm^−1^).^66^ Unless otherwise specified, productions runs were performed in the NPT ensemble using a Nose-Hoover^67^ thermostat and a Parrinello-Rahman barostat^68^ with time constants of 5 ps. In all cases, the target pressure was fixed to 1 atm.

Following a strategy that has been extensively applied to proteins and to other RNA systems in the past,^69–71^ we then used the Hamiltonian replica exchange with solute scaling (REST2) sampling method^72^ for 1 *μ*s over 16 replicas. Briefly, a progressive scaling factor *λ*_*i*_ (spanning from 1 for the first replica to 0.56 for the 16-th replica) is applied on the potential energy of each phase space configuration X of replica “i” (see Eq. 1):

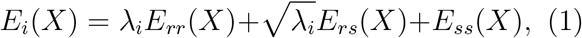

where *E*_*rr*_(*X*) denotes the RNA-RNA interactions of configuration X, *E*_*rs*_(*X*) the RNA-solvent interactions, and *E*_*ss*_(*X*) the solvent-solvent interactions. The scaling factor affects the potential energy of the biomolecule, which allows to mimic the effects of a temperature increase while ensuring a better energetic overlap between adjacent replicas as compared to the regular temperature replica-exchange strategy. Each *λ*_*i*_ can be associated with a corresponding temperature *β*_*i*_ such that *λ*_*i*_ = *β*_*i*_*/β*_*ref*_ . We had shown that an effective temperature 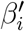 can be attributed to the *i*-th replica evolving on a rescaled potential energy surface with a *λ*_*i*_ scaling factor^18^ such that follows^46^

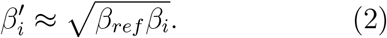

In this work, the corresponding temperatures *T*_*i*_ (associated with the *λ*_*i*_) are distributed along the *N* = 16 replicas spanning from 0 to *N* − 1 according to:

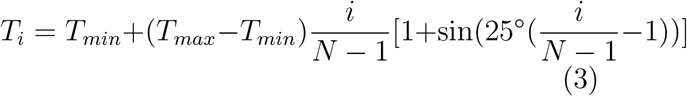

with *T*_*min*_ and *T*_*max*_ being respectively 300 K and 540 K. As compared to usual geometric distributions, this function ensures a denser spacing of replicas at low temperatures.

Every 5 ps, attempts are made to exchange configurations between adjacent replicas in order to increase the conformational sampling, with a success rate around 10–20% along the rescaling factor ladder.

## Data analyses

The fraction of hydrogen-bond (HB) contacts was determined as

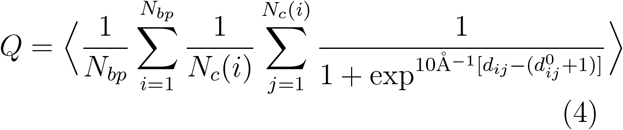

Here, *N*_*bp*_ is the number of possible complementary base-pairs in the duplex, *N*_*c*_(*i*) the number of possible HB contacts in the *i*-th base pair (2 for A-U and 3 for G-C), *d*_*ij*_ the distance (in Å) between the donor hydrogen and the acceptor for each HB *j* involved in the base-pair *i*, and 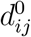 the same distance averaged over the contacts found in experimental homoduplexes. ^46^ *Q* varies between 0 and 1, with 1 being a completely hybrized double-stranded RNA and 0 corresponding to separated single strands.

To obtain the melting curves, we calculated the fraction of bound duplex conformations *f*_*A*_(*T*) defined as those with *Q* larger than 0.1 at a given temperature (we have verified that varying this cut-off did not change the melting curves^46^). In simulations of a single duplex, the melting temperature *T*_*m*_ corresponds to the temperature at which *f*_*A*_(*T*) *<* 2*/*3.^73^ For the 2d distributions of base pairing along the sequence at increasing temperature, we calculated the fraction of formed HBs for a given base-pair using a threshold of 3 Å.

For the artificial sequences lack experimentally-determined melting temperatures, we used the parameters of the latest version of a nearest-neighbor model dedicated to the prediction of the temperature of denaturation of double-stranded RNA^21^ to compare to the results obtained by our simulations.

All trajectories are analyzed once a steady-state is reached, which typically occurs during the first 500 ns of the simulations. ^46^ Therefore, we typically discard the first half of the microsecond-long trajectories and we do analyze the second 500-ns portion.

**Table 2:** Average hydrogen-bond (HB) distances found in the experimental structures of exp-10_GC_ (1RNA) and exp-14_AU_ (4MS9).

|  |  |  |
| --- | --- | --- |
| Uridine-Adenine | O4-H61 | 1.968 Å |
|  | H3-N1 | 1.863 Å |
| Cytosine-Guanine | H42-O6 | 1.908 Å |
|  | N3-H1 | 1.880 Å |
|  | O2-H22 | 1.800 Å |

## Results

### Artificial strands with controlled properties

We first discuss the melting properties of artificial strands with controlled characteristics, i.e., with well-defined G-C/A-U content and ordering, and compare it to the results we had obtained for experimental structures containing either weak or strong base pairs, and artificial strands with random distributions of such pairs.^46^ The exploration of the duplex conformations along the temperature scale is achieved using Hamiltonian replica-exchange (Replica Exchange with Solute Tempering (REST2)^72^) simulations. Indeed, while a more direct approach would be to use replicas in temperature, this is in practice hardly applicable to all-atom simulations in explicit solvent for biomolecules of decent sizes, as the number of involved replicas would rapidly grow to the hundreds.

The central idea is to run several replicas of the system with an increased rescaling of the potential energy terms involving the biomolecule, and to allow for periodic, random exchanges along the rescaling ladder that follow detailed balance criteria. While this is typically exploited to enhance the exploration of the unperturbed replica, by benefiting from configurations generated with rescaled potentials (as already applied for RNA^69–71^), it can also be used to study the biomolecule melting properties by extracting the information from all replicas. ^18,19,47–50^ Indeed, we had shown that we could attribute an effective temperature of the biomolecule^18^ for each of the replica with a rescaled potential energy, and therefore, follow the duplexes conformations explored at increasing temperatures.

### Investigated systems

We first performed replica-exchange simulations for seven duplex structures of identical length (12 residues in each strand, Tab. 3), using the same RNA/water forcefield combination as in our previous work (Amber ff99 forcefield with bsc0 and *χ*_OL3_ corrections^54^ in TIP3P water, see Methods). In order to verify whether an approach based on artificial strands would very much differ from that using experimental sequences and structures, we first selected two duplexes with either G-C base pairs only (art-(CG)_6_), or A-U only (art-(UA)_6_). We then mixed A-U and G-C base pairs to equal proportions, but with defined patterns, either by alterning G-C and A-U base pairs (art-(AG)_6_), by flanking strong base-pairs with weak ones (art-A_3_G_6_A_3_), or the opposite (art-G_3_A_6_G_3_). Finally, we also generated two sequences (art-RND_1_ and art-RND_2_) with a random A-U/G-C content and sequence.

**Table 3:** Artificial double-stranded RNAs. From left to right: abbreviation used in the current work; sequence ; nearest neighbor model^21^ melting temperature at the simulation concentration ; simulation melting temperature (simulation concentration).

| Name | Sequence | Pred. T<sub>m</sub><br>(Sim. CC.) | Sim T<sub>m</sub><br>(Sim. CC.) |
| --- | --- | --- | --- |
| art-(CG)<sub>6</sub> | 5'-(CG)<sub>6</sub>-3' | 359.1 | 372.2 |
| art-(UA)<sub>6</sub> | 5'-(UA)<sub>6</sub>-3' | 323.5 | 323.6 |
| art-(AG)<sub>6</sub> | 5'-(AG)<sub>6</sub>-3' | 347.3 | 356.9 |
| art-A<sub>3</sub>G<sub>6</sub>A<sub>3</sub> | 5'-A<sub>3</sub>G<sub>6</sub>A<sub>3</sub>-3' | 352.2 | 374.1 |
| art-G<sub>3</sub>A<sub>6</sub>G<sub>3</sub> | 5'-G<sub>3</sub>A<sub>6</sub>G<sub>3</sub>-3' | 350.3 | 357.4 |
| art-RND<sub>1</sub> | 5'-UAGGAGGCUAGA-3' | 351.5 | 364.4 |
| art-RND<sub>2</sub> | 5'-GAACUGUAUGAU-3' | 338.6 | 348.0 |

Experimental melting data is not available for these artificial sequences. We therefore use nearest neighbor models to estimate the melting temperatures 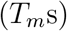. In Tab. 3, we report the melted temperatures expected with a state-of-the-art, recent instance of a nearest neighbor model.^21^ These values correspond to a strand concentration equal to that of the simulation (note that the ion concentrations are different in our simulations as compared to these models, but as discussed in the Methods section, this has a very limited impact on the melting temperatures). Not surprisingly, the 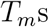 are lower for the strands with weak base pairs and stronger for the strands containing GC base pairs. All the other sequences contain a mixture of G-C and A-U base pairs, leading to pretty similar melting temperatures for those containing the two types of base-pairs in similar proportions, except art-RND_2_ which is rich in A-U base pairs and thus has a lower melting temperature as compared to the other mixed sequences.

### Simulated melting curves and melting temperatures

We discuss the results of microsecond-long REST2 simulations (see Methods), which are analyzed once a steady-state is reached. In order to track the duplex separation, we use a collective variable (CV) that quantifies the normalized, smoothed fraction of hydrogen-bonds (HBs) formed between the two complementary strands, which varies between 0 (fully separated strands) and 1 (fully formed duplex), see Methods for definition and technical details.

In Fig. 1A, we show the ensemble-averaged ⟨*Q*⟩ as a function of temperature for all seven systems. As temperature increases, ⟨*Q*⟩ typically drops from a value close to 1 at ambient temperature to 0 at the highest investigated temperature, illustrating the progressive melting of the duplexes as temperature increases. However, typical optical melting curves determined from experimental measurements do not report on ⟨*Q*⟩, but on the fraction of formed duplexes. As discussed in the Methods section, we define a duplex as a state for which at least a small fraction of HBs (which exact value has only a very small impact on the estimated melting temperature) are formed. From the distributions of *Q* at each temperature, we can thus determine the fraction of associated duplexes *f*_*A*_, and therefore determine the melting curves shown in Fig. 1B. Finally, *T*_*m*_ corresponds to the temperature for which *f*_*A*_(*T*) = 2*/*3 (which is not exactly 1/2 as in experimental bulk measurements, because of differences in the fluctuations of the local duplex concentration, as shown in ref.^73^).

**Figure 1:**
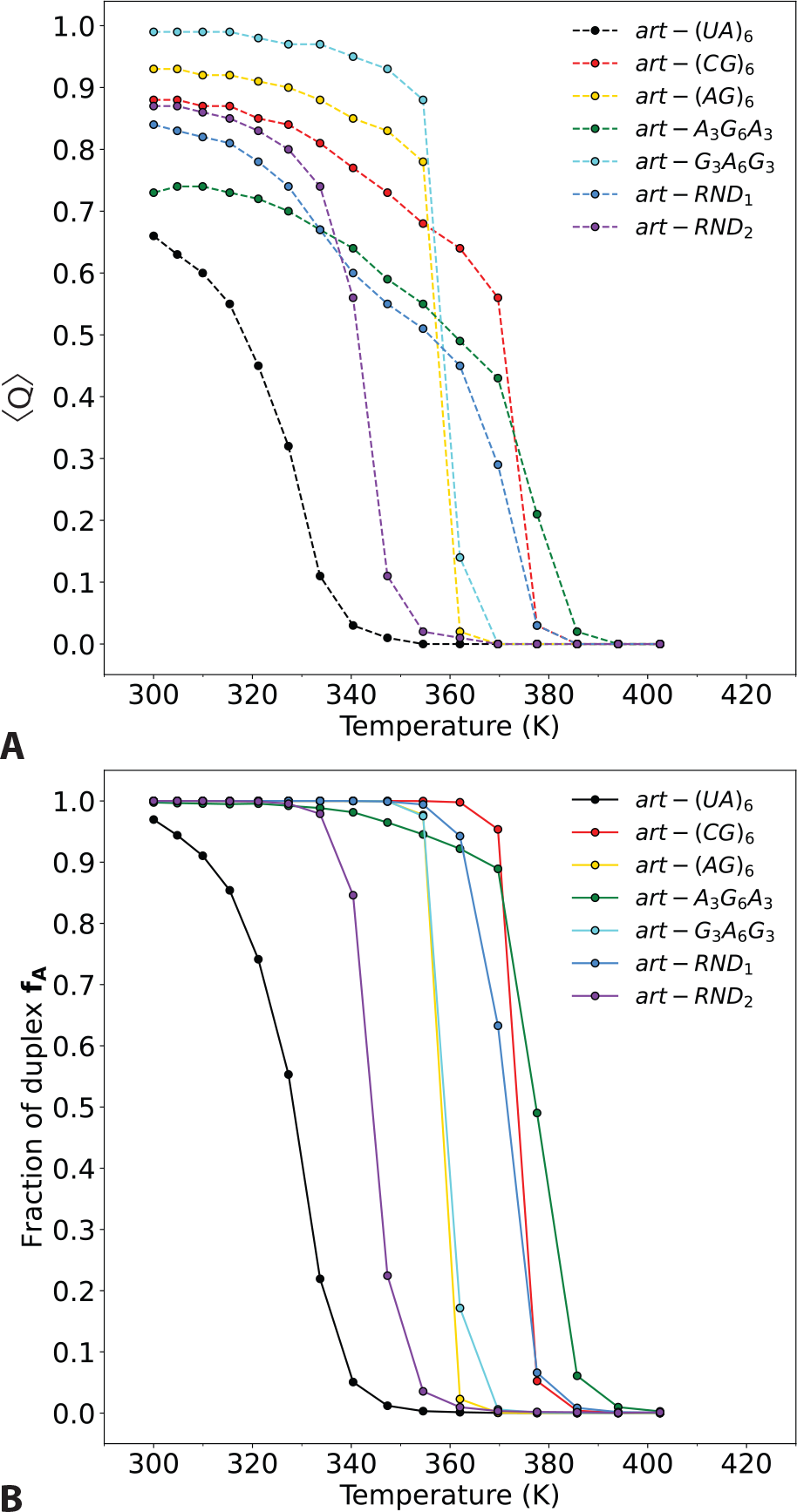
Denaturation of duplexes with controlled or random sequences. (A) Ensemble average of the fraction of native HB contacts (Eq. 4) at increasing temperatures. (B) Fraction of associated duplex conformations *f*_*A*_ at increasing temperatures. The 2/3 limit that allows for the determination of the melting temperature is indicated in gray.

### Comparison with nearest-neighbor models

We can now compare the simulated *T*_*m*_ to the expectations from the model, as shown in Fig. 2A for all duplexes. These results confirm the good agreement between simulated and expected *T*_*m*_ for all types of duplexes. In particular, simulations can very well reproduce the gradual increase in the melting temperature with the fraction of G-C base-pair in the sequence, as observed before. Besides force-field limitations, which will be addressed in a subsequent section, we note that such a good agreement between the simulated and predicted melting temperatures is remarkable but hides some limitations of our approach. The exploration of the duplexes conformational space, and notably in their dissociated geometries, may be imperfect, and the determination of the melting temperatures results from an a posteriori reconstruction of a temperature scale that is not the physical temperature of the system. However, a critical aspect of our approach is that it can capture the sequence-dependence of the duplexes melting temperature, and how this later increases with the G-C content of the duplex.

**Figure 2:**
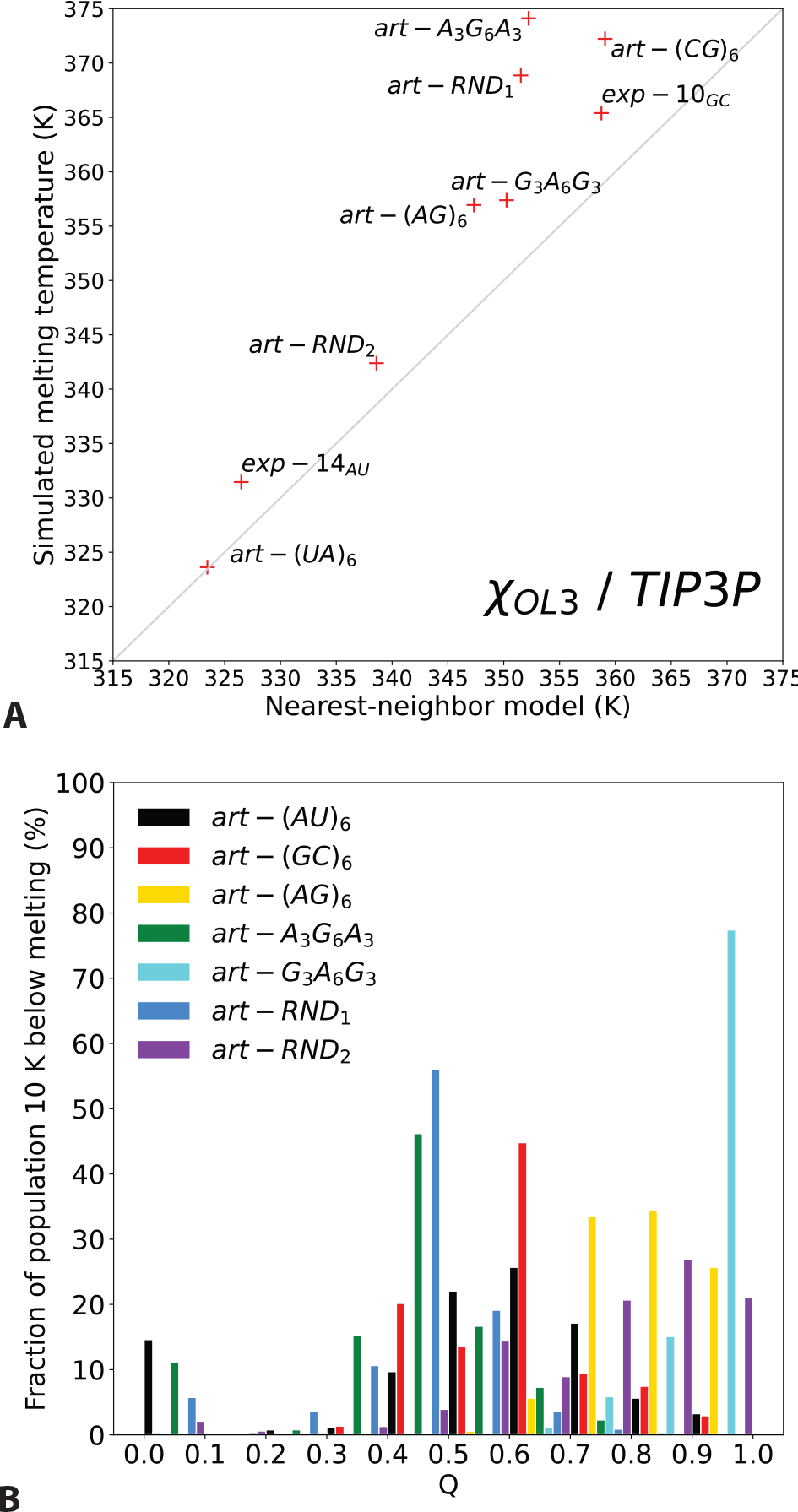
Comparison with nearest-neighbor models and evidence of partially-melted intermediates. (A) Correlation between the simulated melting temperatures and the melting temperature estimated from a state-of-the-art nearest-neighbor model. ^21^ (B) Distributions of the fraction of HB contacts in the replica located ≈ 10 K below melting.

### Melting cooperativity

We now more specifically discuss the shape of the curves representing the average fraction of formed base-pairs as a function of temperature (Fig. 1A), which readily points toward some peculiar behavior of several artificial sequences, and which cannot be predicted from nearest-neighbor models. Despite differences in the average number of contacts at ambient conditions, the melting behavior of a first set of duplexes exhibit some clear, gradual melting with sigmoidal decay toward 0 as temperature increases. These include the artificial homoduplexes (in that sense that they only contain either strong or weak base-pairs), art-(CG)_6_ and art-(UA)_6_, as well as that of the strands with random sequences (art-RND_1_ and art-RND_2_). As already observed before,^46^ a two-state behavior is not observed in the distribution of *Q* just before melting (Fig. 2B). These distributions show peaks with intermediate values of *Q*, that are very different from the expectations of a 2-state model, that would result in a bimodal distribution, with two peaks centered around *Q* ≈ 0 and *Q* ≈ 1.

Two duplexes exhibit a peculiar behavior. When the core of the duplex is made of weak A-U base pairs, and the extremities of strong G-C base pairs, the melting is much more abrupt; within a few K, the duplex goes from completely formed to completely separated (Fig. 1A and B). In the distributions, this is evidenced by the absence of structures with intermediate values of *Q* close to melting (Fig. 2B and Fig. SI1 and 2 for a more complete temperature dependence of these distributions); this duplex is perhaps closest to what would be a two-state behavior. In contrast, now flanking a strong G-C core with weak A-U extremities, results in a denaturation that is much more spread in temperature as compared to the other systems (Fig. 1A). This is even more prominent in the distributions along *Q*, that are very wide (Fig. 2B). On average, almost 50% of HB interactions lost 10 K before melting, which is consistent with the very progressive loss of structure seen in the melting curves. These observations suggest that the specific base-pair ordering can tune the cooperativity of the duplex formation/denaturation, at fixed G-C/A-U content and despite pretty similar melting temperature as predicted by nearest neighbor models. As already discussed before,^46^ the observed intermediates do not seem to be artefacts of the employed enhanced sampling strategies, as they are stable when propagated at real physical temperatures (Fig. SI3).

### Local melting

We now have a closer look at the individual melting of each base-pair interaction along the sequence as temperature increases. We determine the respective populations of associated and dissociated states for each base-pair, resulting in 2-dimensional histograms (Fig. 3 and SI4 for other sequences), which provide a molecular picture of the three types of behavior identified above.

**Figure 3:**
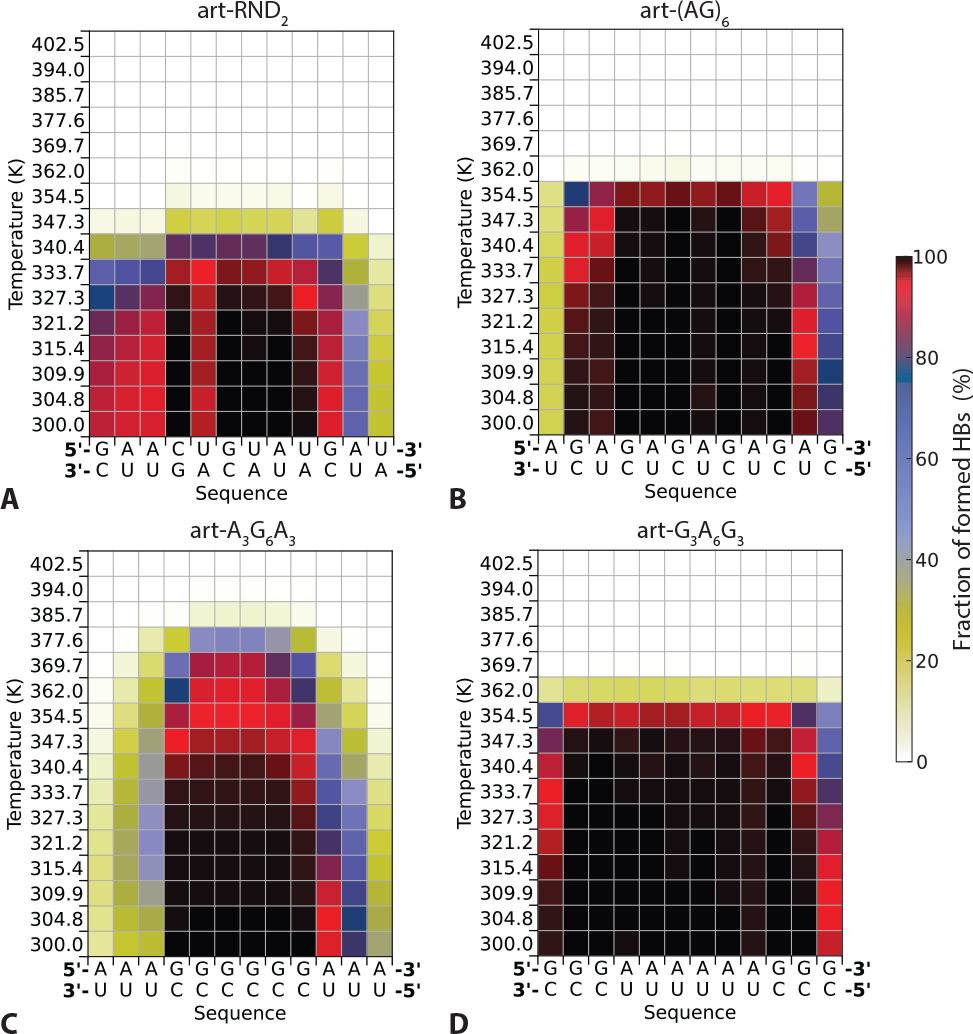
Local duplex melting. Fraction of formed base-pairs (color scale) between the two complementary strands of (A) art-RND_2_, (B) art-(AG)_6_, (C) art-A_3_G_6_A_3_ and (D) art-G_3_A_6_G_3_ along the sequence (x axis) and at increasing temperature (y axis).

For most structures (pure G-C, pure A-U, random sequences and alternating mixture of A-U and G-C base pairs, see e.g. Fig. 3A and B), the denaturation is progressive and the extremities are destabilized as temperature increases, with significant base fraying spanning up to 3 residues at each extremity before full denaturation occurs, similar to what had been observed for experimental duplex structures before.^46^

The denaturation of art-A_3_G_6_A_3_, which was observed to occur on a wide temperature range, is in fact occurring in two stages (Fig. 3C): first, the weak extremities, already poorly stable at ambient temperature, are readily denatured around 330-340 K, i.e., the melting temperature for A-U duplexes. At this temperature, the core G-C structure begins to unfold by its extremities, with G-C base fraying propagating toward the duplex center before full separation occurs around 370 K, i.e., the melting temperature of a pure G-C duplex. Overall, conformational changes occur along a 60 K temperature scale due to this two-stage denaturation.

Another extreme is provided by the art-G_3_A_6_G_3_ duplex (Fig. 3D). Base fraying is similar to what is observed for a pure G-C duplex, with mainly 1 base-pair at each extremity that is opening around 340 K. But, because this temperature now exceeds the melting temperature of the A-U core sequence block, the propagation of temperature induced destabilization through the extremities leads to the global and sudden duplex denaturation. For this duplex, the local melting distribution therefore explains the two-state behavior inferred from the melting curves and the distributions of the number of HB interactions (Fig. 2B).

### Hairpin and circular motifs

Since the duplex extremities appear as weak points of the thermal unfolding, we now discuss the melting behavior or artificial constructs that consist in some of the sequences already simulated where either one, or both extremities, are artificially maintained in a closed state regardless of the temperature. This is obtained by adding local upper wall potentials to terminal base-pair interactions. While such strategy results in artificial hairpin and circular RNA conformations that have noting to do with experimentally-relevant objects, they allow to specifically target the effects of base-fraying on the thermal stability of dsRNA, independently of e.g. the degrees of freedom of loop motifs in more relevant hairpin motifs.

In Fig. 4, we show the 2-dimensional melting histograms showing the propagation of unfolding along the sequence as temperature increases, for art-(UA)_6_ (Fig. 4A-C) and art-(CG)_6_ (Fig. 4D-F). They evidence three important consequences of forbidding base-fraying at one or two extremities. Not surprisingly, the first important effect is to shift melting temperatures towards higher values, typically by 10– 15 K when both extremities are blocked. This is a consequence of the fact that at a given temperature, the fraction of formed HBs for base-pairs at the vicinity of the blocked extremity are much higher compared the free duplex case. The effect is quite spectacular for art-(UA)_6_, whose HBs are weakly stable even at ambient temperature (Fig. 4A), but become much more compact with joint extremities (Fig. 4B-C). A second important observation is that such effect propagates along more than half the length of these dodecamers. typically along 8-9 residues. Quite strikingly, the base-pair that is 9 residues away from the only blocked extremity gets stabilized even when it is only 3 residue away from the other, free end of the duplex.

**Figure 4:**
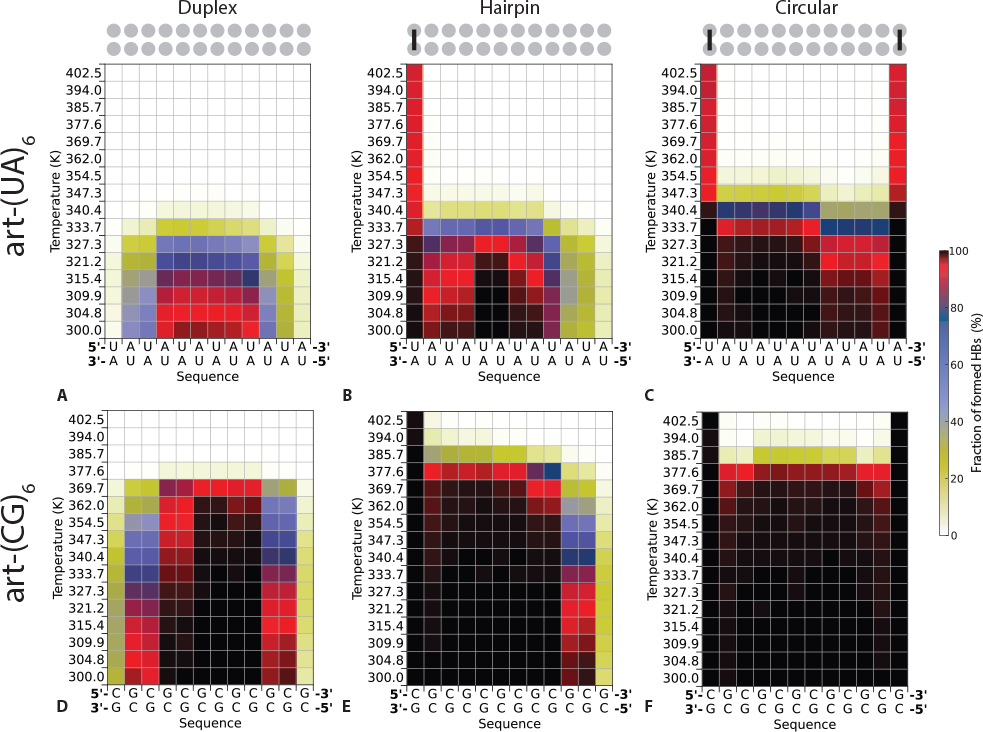
Local duplex melting with constrained extremities. Fraction of formed basepairs (color scale) between the two complementary strands of art-(UA)_6_ with (A) free extremities, (B) with one blocked end (hairpin state) and (C) two blocked end (circular RNA). (D), (E) and (F): similar data for art-(CG)_6_.

The final important consequence of impeached fraying is that thermal unfolding becomes much more cooperative and close to a two-state behavior, especially for art-(CG)_6_ (Fig. 4F). In particular, no intermediate states are observed close to melting and there is a sharp transition between a situation where the duplex is fully formed, with no significant deformation of the duplex structure before the melting temperature.

### Forcefield dependence

We finally check whether our conclusions regarding the separation mechanism of RNA duplexes are robust against the specific choice of the RNA and solvent forcefields, which was only briefly alluded to before,^46^ and we now discuss it in a more systematic way.

We first tested on the same set of artificial duplex sequences, together with the two experimentally-available structures studied before^46^ (exp-14_AU_ ^74^ and exp-10_GC_ ^75^) more recent, versatile RNA forcefield that include additional specific reparametrizations of backbone dihedral angles (*ϵζ*_OL1_-*β*_OL1_ corrections^58,59^), together with the TIP3P water model. As shown in Fig. 5, including the first of these corrections, or both, results in a decreased stability for all investigated systems, for which the simulated melting temperature are typically lowered by 20–30 K as compared to our original simulations (Fig. 2A). Overall, the stability trend, i.e., the increase of the melting temperature with the G-C content is still observed, suggesting that this forcefield captures the basic ingredients of the sequence-dependence of the melting temperature. However, large deviations are noticed for some duplexes, for example with sequences that have very similar predicted 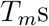 but very different simulated ones (Fig. 5).

**Figure 5:**
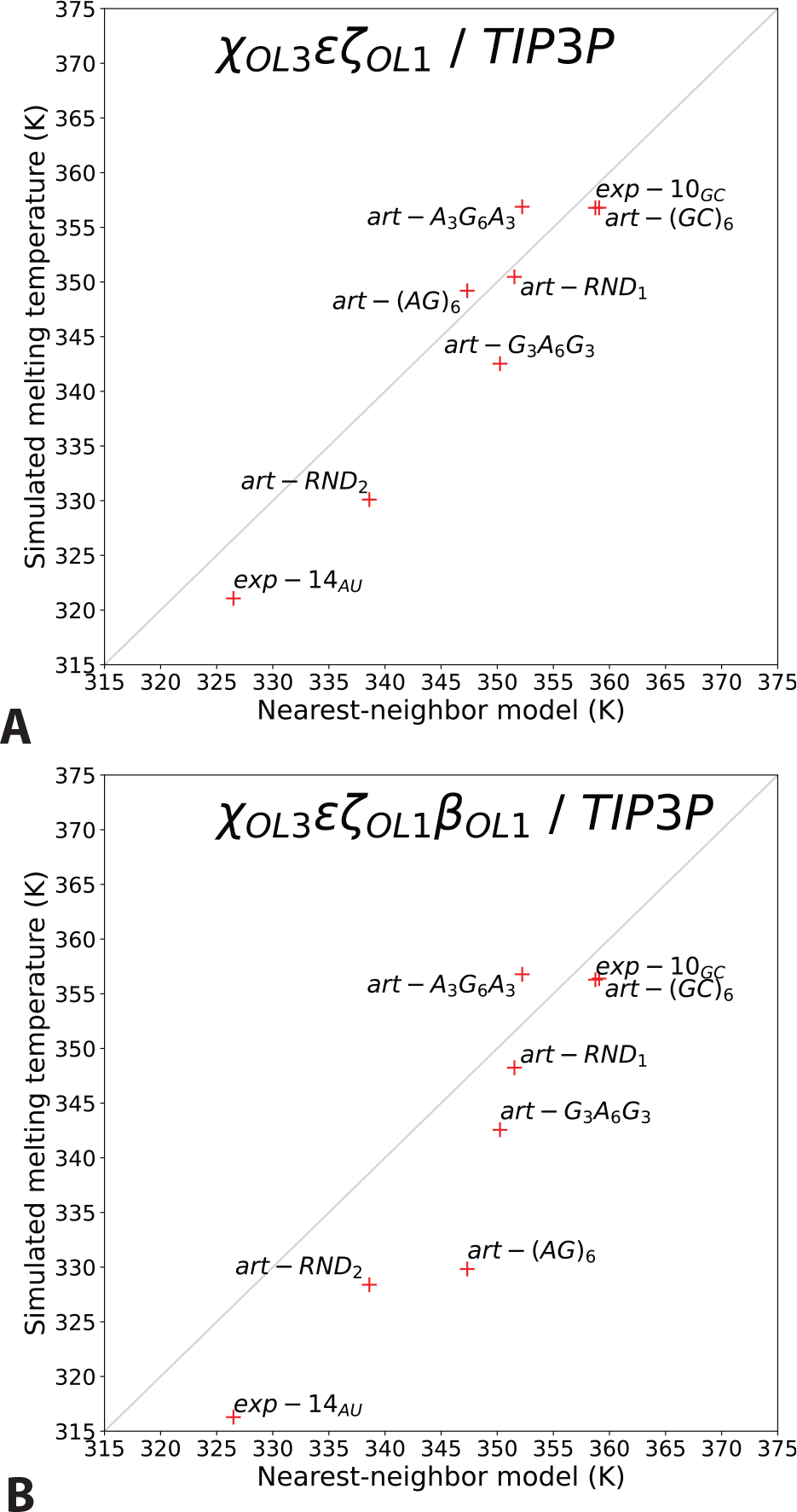
Correlation between the simulated melting temperatures and the melting temperature estimated from a state-of-the-art nearestneighbor model^21^ for the RNA 99sb/bsc0/*χ*_OL3_-*ϵ ζ*_OL1_ forcefield in TIP3P water (A) and the RNA 99sb/bsc0/*χ*_OL3_-*ϵ ζ*_OL1_-*β*_OL1_ forcefield in TIP3P water (B). Note that a simulation melting temperature could not be determined for art-(UA)_6_, which were already below the folded fraction threshold in the unperturbed replica at 300 K. The values of the melting temperature used for this figure cab be found in Table SI1.

The limitations of this forcefield are further illustrated by the very poor stability of duplexes containing only weak base-pair interactions. For these two systems (art-(UA)_6_ and exp-14_AU_), we could not determine a melting temperature as the duplex was readily not stable at 300 K. For the two experimental structures, the duplexes simulated with the 99sb/bsc0/*χ*_OL3_-*ϵ ζ*_OL1_-*β*_OL1_ forcefield are much less stable as compared to the original 99sb/bsc0/*χ*_OL3_ version, which results for exp-14_AU_, in a folded fraction that is below 50 % under ambient conditions. We note that while the melting curves are sensitive to reconstruction of the effective temperature scale, the data at 300 K, i.e. in the unperturbed replica, is devoid of any methodological artefacts. These observations are at odds with the experiments,^74^ suggesting that the 99sb/bsc0/*χ*_OL3_-*ϵ ζ*_OL1_-*β*_OL1_ forcefield might not be adequate to study RNA duplex systems.

In order to more extensively explore other forcefields for RNA and for the water solvent, we selected one of the random sequences and repeated our simulations by testing the following combinations (Fig. 6). First, we varied the water model only, and used the SPC/E^57^ or the OPC^57^ water models instead of TIP3P.^56^ These had a rather limited impact on the duplex thermal stability, although the melting temperature was slightly increased with SPC/E (Fig. 6A). As mentioned above, the recent corrections of the original RNA forcefield (*ϵ ζ*_OL1_-*β*_OL1_ corrections^58,59^), in TIP3P water, result in much weaker interstrand interactions and thus, a duplex that has a rather surprisingly-low melting temperature. On the other hand, the very recent gHBfix21 potentials^60^ with the original *χ*_OL3_ forcefield^54^ including vdW phosphate corrections,^61,62^ in OPC,^63^ results in a very stable structure, with a melting temperature that shifts by about 25 K compared to *χ*_OL3_ (Fig. 6A). But importantly, the gradual unfolding mechanism by the propagation of base fraying toward the inner part of the duplex as temperature increases is seen for all models, typically spanning 3-4 residues on each extremity before full separation is observed (Fig. 6B). As a consequence, the molecular mechanism of the duplex dehybridization appears to be quite insensitive to the specific choice of forcefield of the solvent and the RNA molecules.

**Figure 6:**
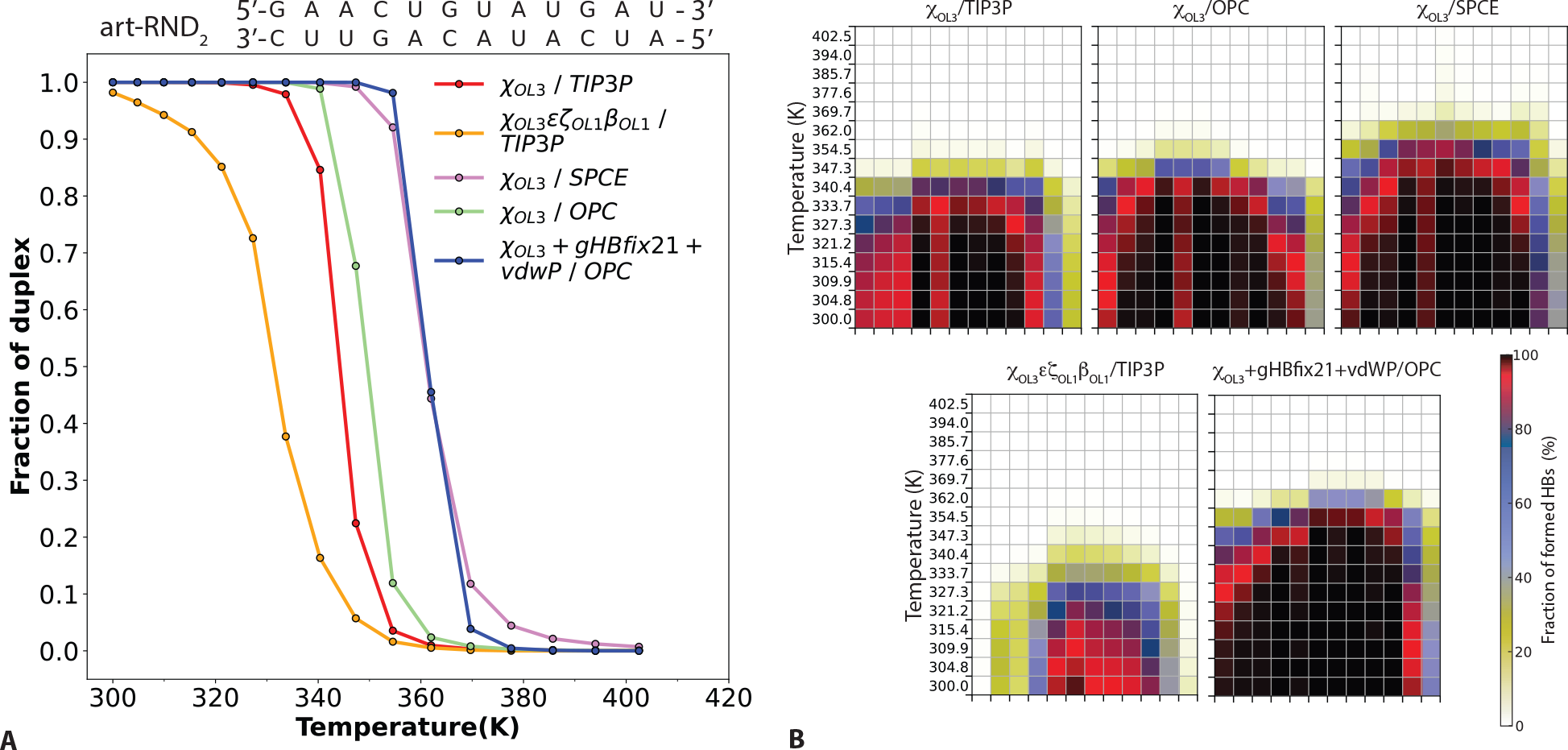
Forcefield dependence of melting properties. (A) Fraction of associated duplex conformations *f*_*A*_ at increasing temperatures for the art-RND_2_ duplex using five different forcefield combinations. (B) Local duplex melting: fraction of formed base-pairs (color scale) between the two complementary strands of art-RND_2_ for these five set of parameters.

## Discussion and Conclusions

In a previous work, we had shown that an all-atom molecular dynamics simulation strategy relying on an Hamiltonian replica exchange scheme allowed to scan the duplexes’ conformational landscapes at several increasing temperatures at the same time, and at a reasonable computational cost.^46^ This approach is both able to reproduce the experimental trends measured for the duplexes thermal stability, and at the same time, enables to provide relevant mechanistic details about their denaturation when approaching the melting temperature. Most importantly, we had shown that around melting, the duplexes are considerably distorted, with base-fraying at their extremities typically spanning at least 2-3 base pairs. This suggests that a 2-state model provides only a simplistic picture of the RNA duplex melting, as no significant population of fully-formed duplexes is observed around melting, in contrast to the expectations of a model considering a temperature-dependent shift in populations of two well-defined thermodynamic basins (one fully formed duplex and the separated strands). This confirmed previous evidence for complex kinetic and mechanistic (de)hybridization pathways, long-lived intermediates, and extensive base-fraying at increasing temperatures,^15,20–24,34,35,39,40^ none of which are expected would a 2-state model be valid.

Although very insightful, this previous study was limited to a few duplex structures, and focused exclusively on one forcefield, although preliminary tests indicated that the molecular picture provided by these simulations was forcefield independent.^46^ The current contribution brings at least three important novelties. First, we investigate the sequence dependence of the dehybridization mechanism by studying artificial strands with controlled sequences, and show that our conclusions remain valid and applicable for all investigated duplexes, but with interesting dependences upon the specific ordering of weak and strong base-pairs among the sequence. Second, as the weak points for thermal unfolding of duplexes are their extremities, we study artificial constructs in which one or two duplex extremities are locked. Finally, we give ample evidence that a variety of forcefield combinations all provide the same mechanistic picture, but at the same time demonstrate that some forcefield combinations result in very unstable duplex conformations even at room temperature, at odds with the experiments. We now discuss each of these aspects.

We choose to focus on a variety of artificial dodecamer duplex constructs, containing: either G-C, or A-U, base pairs only; a mix of A-U and G-C base pairs in equal proportions, but with defined patterns, either by alterning G-C and A-U base pairs, by flanking strong base-pairs with weak ones, or the opposite (art-G_3_A_6_G_3_); and two duplexes with a random A-U/G-C content and sequence. The simulated melting temperatures are in excellent agreement with the expectations of state-of-the-art nearest-neighbor models.^21^ Crucially, our approach is able to precisely account for the sequence-dependence of the melting temperatures, a conclusion already reached in our previous study but on a very limited number of duplexes. Now extended to a total of 9 duplexes with size ranging from 10 to 14 base-pairs, this suggests that our simulation strategy offers a reliable account of the duplexes thermal stability. This legitimates the use of these simulations to provide a molecular picture of the thermally-induced strand separation, which cannot be provided by nearest-neighbor models and which is not easily accessible in the experiments.

A detailed investigation of the position-dependent Watson-Crick formed inside the duplexes reveals that these partially-melted structures correspond to significantly-frayed structures (summarized in Fig. 7). For most types of sequences (A-U only, G-C only, alternating G-C and A-U base pairs), 2-3 base pairs at each extremity of the dodecamer duplexes are significantly distorted and partly broken close to melting, supporting the interpretation of previous experimental measurements.^34^ Two specific constructs exhibit more extreme behavior. For a duplex consisting of a strongly-bound G-C core and AU extremities, the separation of the A-U base pairs clearly occurs before that of the G-C core. This leads a very wide melting curve, and to the presence of a very distorted duplex with a strong core and melted extremities. The mirror scenario (A-U core with G-C extremities) is completely different, and is here the only system with an apparent 2-state behavior. Indeed, once the much stronger G-C base-pairs (as compared to A-U) at the extremities begin to melt, the entire duplex falls apart because of the weak interactions in its core. All these observations are in agreement with recent temperature-jump infrared spectroscopy experiments, that were interpreted using a simple lattice model predicting the same behavior.^34^ Here, we show that the underlying atomistic details can be provided by state-of-the-art molecular simulations.

**Figure 7:**
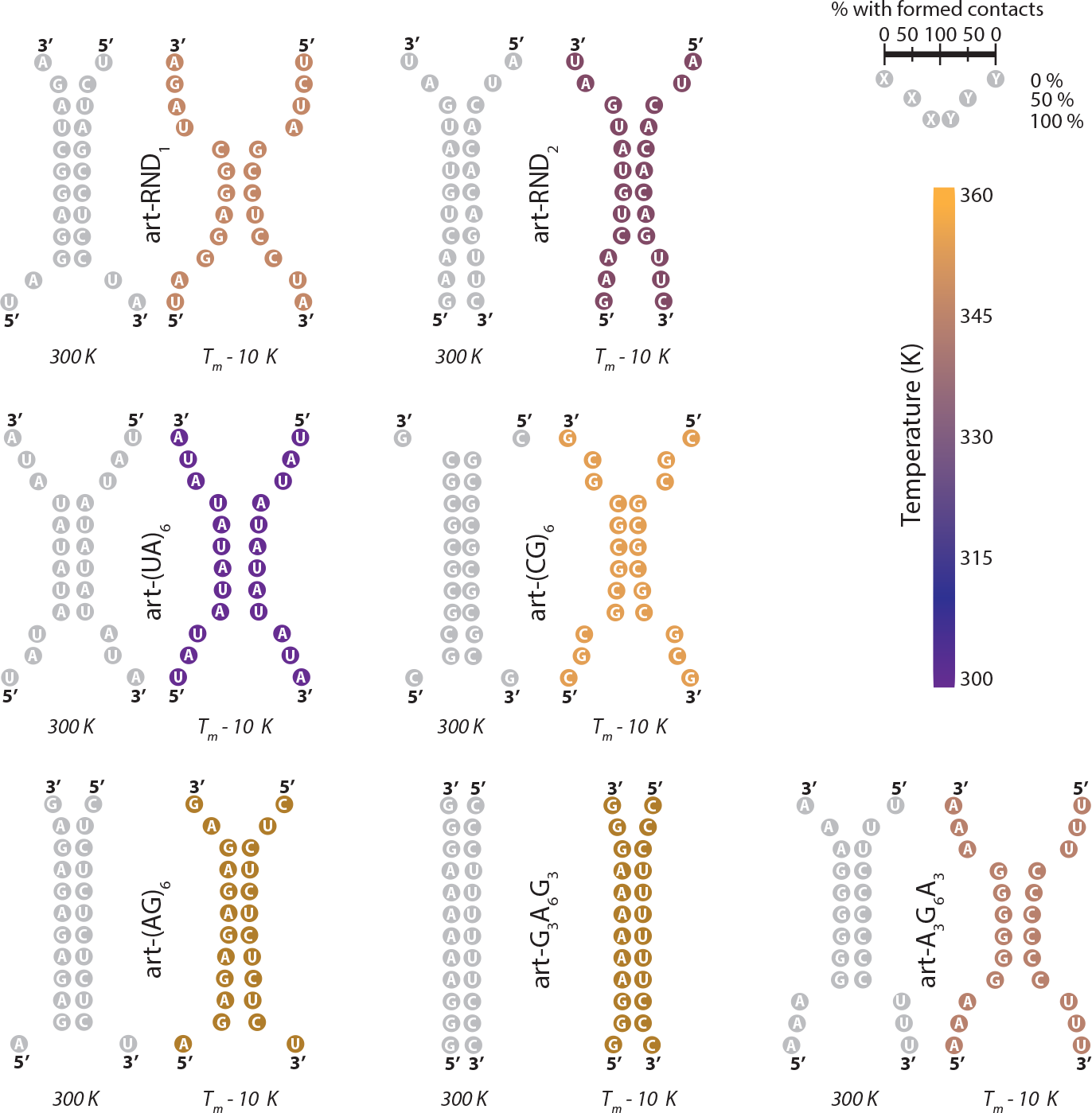
Schematic representation of the fraction of formed base-pairs at ambient temperature (300 K, gray) and 10 K below the melting of each sequence (color scale). In these plots, the distance between two complementary bases is proportional to the fraction of structures that do not base-pair at each temperature (scale on the right hand side). The sequences are organized into 4 families (from top to bottom: experimental structures, artificial structures with only weak or only strong base-pairs, artificial structures with 50/50 and organized sequences, and artificial structures with random sequences).

As unfolding always occurs from the duplex extremities, which appears as the weak points for thermal denaturation, we repeated our simulations for the two duplexes containing one type of canonical base-pairs only, and locking one or both extremities. Not surprisingly, this results in increased thermal stability (typically by 10–15 K), and overall provides much greater occurence of base-pairing interactions, especially for the weak A-U duplexes. This effect is long-range, typically spanning up to 8/9 residues away from the locked extremity, even when located not far from the free end of the formed artificial hairpin. A key consequence of impeached base fraying at both extremities is that unfolding becomes much more cooperative and thus close to the expectations of a two-state model: intermediate, partially-unfolded structures become non-existant around melting.

Our study was originally based on a widely used and versatile RNA/solvent forcefield combination based on the amber 99sb/bsc0/*χ*_OL3_ model for RNA and the TIP3P water models. Although robust, these models can be criticized in that sense that many improvements have been suggested regarding the employed water model or the reparametrization of key RNA torsions or interactions.^25^ While it seems impossible to test the wide range of possible forcefield combinations, we chose to focus on a few key examples to demonstrate that our main conclurions are insensitive to the employed models, but also to point at limitations is some recent but widely used models.

Not surprisingly, changing either the solvent forcefield, the RNA forcefield, or both, results in shifts in the melting temperature of a selected duplex. For example, SPC/E seems to stabilize the structure as compared to TIP3P. The effect is even more pronounced with the very recent gHBfix21 potentials^60^ with the original *χ*_OL3_ forcefield^54^ including vdW phosphate corrections,^61,62^ in OPC.^63^ As compared to the original *χ*_OL3_ forcefield in TIP3P water, the gain in stability exceeds 25 K. While it is hard to conclude whether this is more in line with experimental values or not, considering the approximations of our effective temperature reconstruction method from the Hamiltonian replica exchange simulations, and the fact that the solvent is in fact not “heated” in this approach, this may suggest that these parameters could overstabilize the interactions between the two strands. On the other hand, recent corrections of the original RNA forcefield (*ϵ ζ*_OL1_-*β*_OL1_ corrections^58,59^), in TIP3P water, result in a much lower duplex stability. In particular, the data in the unperturbed replica suggest that the duplex structure, is readily very distorted and not fully formed at 300 K, which is not compatible with experimental observations. This forcefield, although more recent and quite widely used, may not be the best choice for RNA duplexes. Despite these differences in the duplex thermal stability, all investigated forcefield combinations (5 in total) all provide the same molecular picture of the duplex unfolding mechanism, whereby extensive base fraying is observed, and whereby the fully formed duplex does not exist close to melting.

This study enrichs our initial investigation^46^ and confirms that our simulation strategy is a valuable tool complementing previous simulation and experimental approaches. It can provide important molecular details about the mechanisms of RNA duplexes melting with temperature. In particular, we show that the unfolding mechanism of most duplex structures involve partially melted intermediates close to melting, questioning a 2-state thermodynamical model. However, sequences with either strong base-pairs flanked with weak ones, or the opposite situation, give rise to a situation where unfolding is much closer to a 2-state picture in the former case, but even more gradual, 3-state-like, in the latter case. Such results could help design strands with tailored association/dissociation characteristics. Additionally, simulations on equivalent hairpin-like or circular RNA-like structures confirm that the duplex extremities are the weak spots for thermal unfolding, and a thorough study of the forcefield-dependence of these results suggest that our conclusions are very robust and are not sensitive upon the specific choice of the employed molecular models.

## Funding

The research leading to these results has received funding from the European Research Council under the European Union’s Eighth Framework Program (H2020/2014-2020)/ERC Grant Agreement No. 757111 (G.S.). This work was also supported by the “Initiative d’Excellence” program from the French State (Grant “DYNAMO”, ANR-11-LABX-0011-01 to GS).

## Acknowledgments

The simulations presented here benefited from a local computing platform administered by G. Letessier, and benefited from the HPC resources of TGCC under the allocation A0070811005 made by GENCI (Grand Equipement National de Calcul Intensif).

## Conflict of interest statement

None declared.

## Supplementary Information

**Figure 1:**
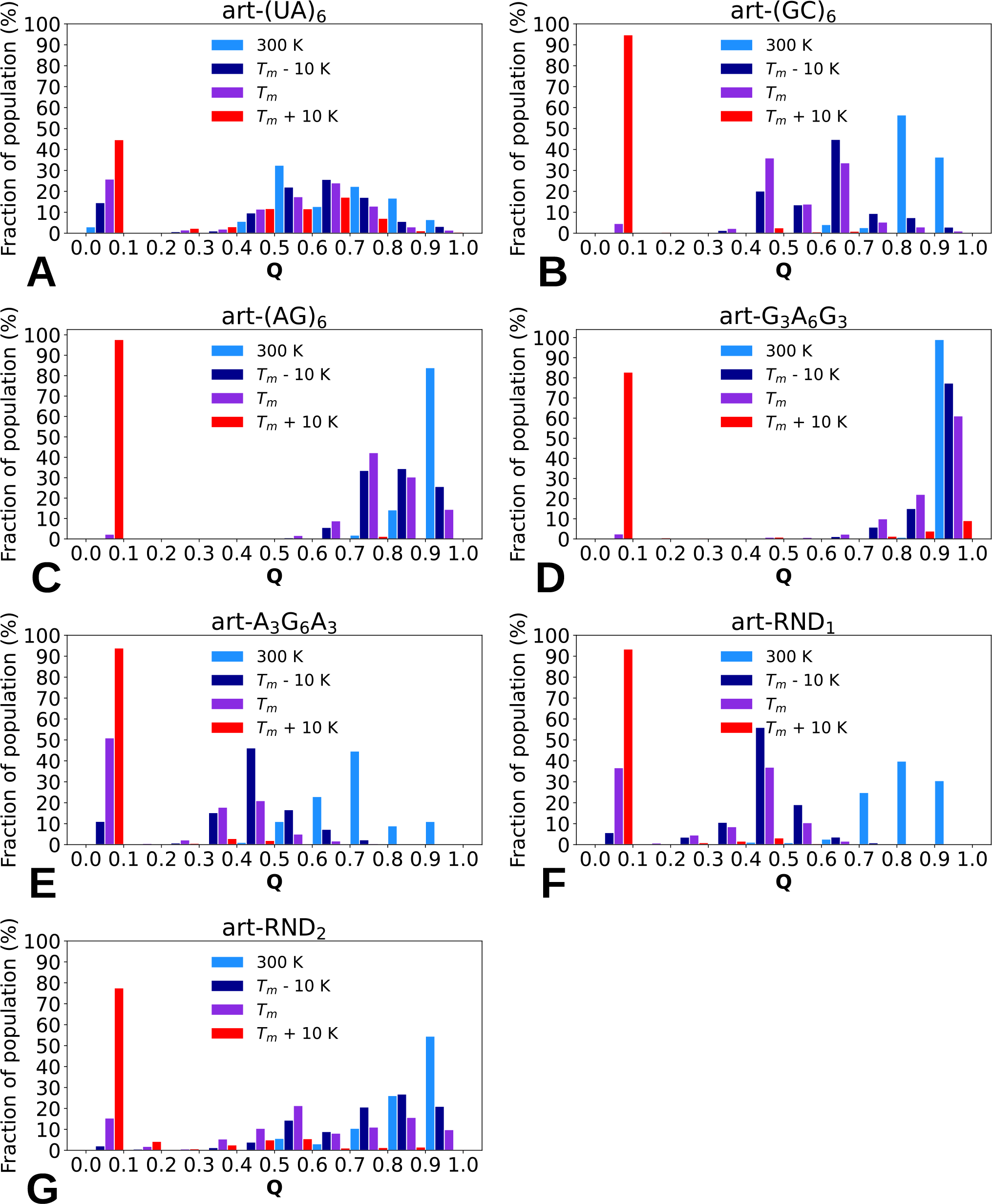
Partially separated intermediates. Distributions of the fraction of HB contacts observed at 300 K (light blue), ≈ 10 K below melting (purple), at melting (magenta) and ≈ 10 K above melting (red), for the artificial strands.

**Figure 2:**
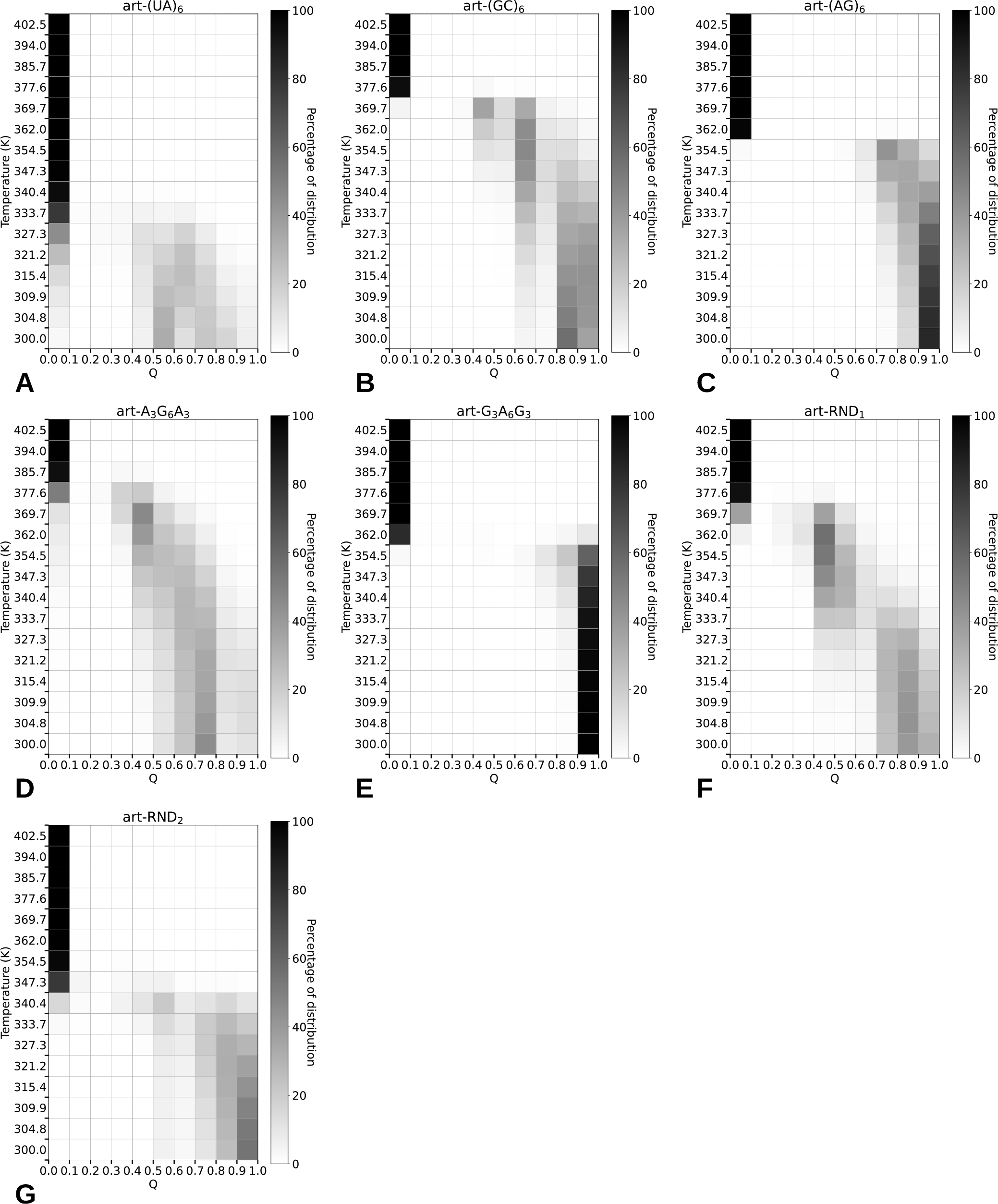
2D maps of duplex melting (fraction of formed HBs, gray scale) for the artificial strands.

**Figure 3:**
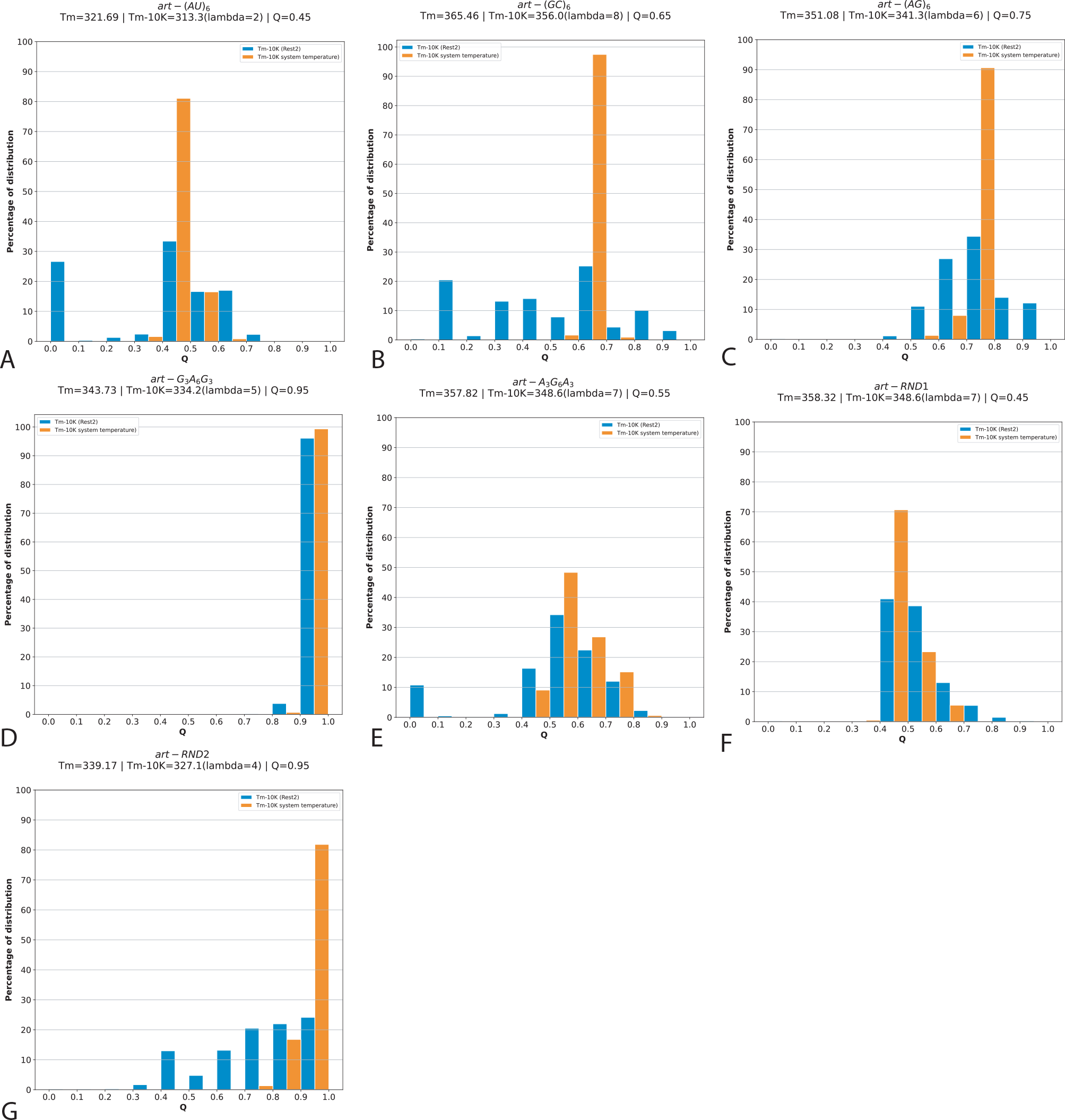
Comparison between REST2 simulations and regular, brute-force MD simulations at similar temperatures. For each system, we show the distributions of *Q* values in blue for the replica that is about 10 K below the effective melting temperature (indicated in the top). We then randomly selected one duplex conformations in the most populated decile (Q values indicated in the upper right corners), resolvated it, and propagated it for 1 *μ*s at a physical temperature corresponding the REST2 replica effective temperature, without any potential energy rescaling, resulting in the distributions shown in orange. These plots suggest that the distorted structures observed around melting are not artefacts of the REST2 setup (as they do not shift, but also that REST2 allows for a much better exploration of the conformational space.

**Table 1:** Table of all effective melting temperatures with the forcefield used during the REST2 simulations and the predicted melting temperatures used for comparisons.

| | NN model | $\chi_{OL3}$ | $\chi_{OL3}\epsilon\zeta_{OL1}$ | $\chi_{OL3}\epsilon\zeta_{OL1}\beta_{OL1}$ |
| --- | --- | --- | --- | --- |
| *exp* – 14<sub>AU</sub> | 326.5 | 323.3 | 321.1 | 316.3 |
| *exp* – 10<sub>GC</sub> | 358.8 | 370.3 | 356.8 | 356.3 |
| *art* – (AU)<sub>6</sub> | 323.5 | 323.6 | 307.7 | / |
| *art* – (GC)<sub>6</sub> | 359.1 | 372.2 | 356.8 | 356.4 |
| *art* – RND<sub>1</sub> | 351.5 | 364.4 | 350.5 | 348.2 |
| *art* – RND<sub>2</sub> | 338.6 | 348.0 | 330.1 | 328.4 |
| *art* – A<sub>3</sub>G<sub>6</sub>A<sub>3</sub> | 352.2 | 374.1 | 356.9 | 356.8 |
| *art* – (AG)<sub>6</sub> | 347.3 | 356.9 | 349.2 | 329.8 |
| *art* – G<sub>3</sub>A<sub>6</sub>G<sub>3</sub> | 350.3 | 357.4 | 342.5 | 342.6 |

**Figure 4:**
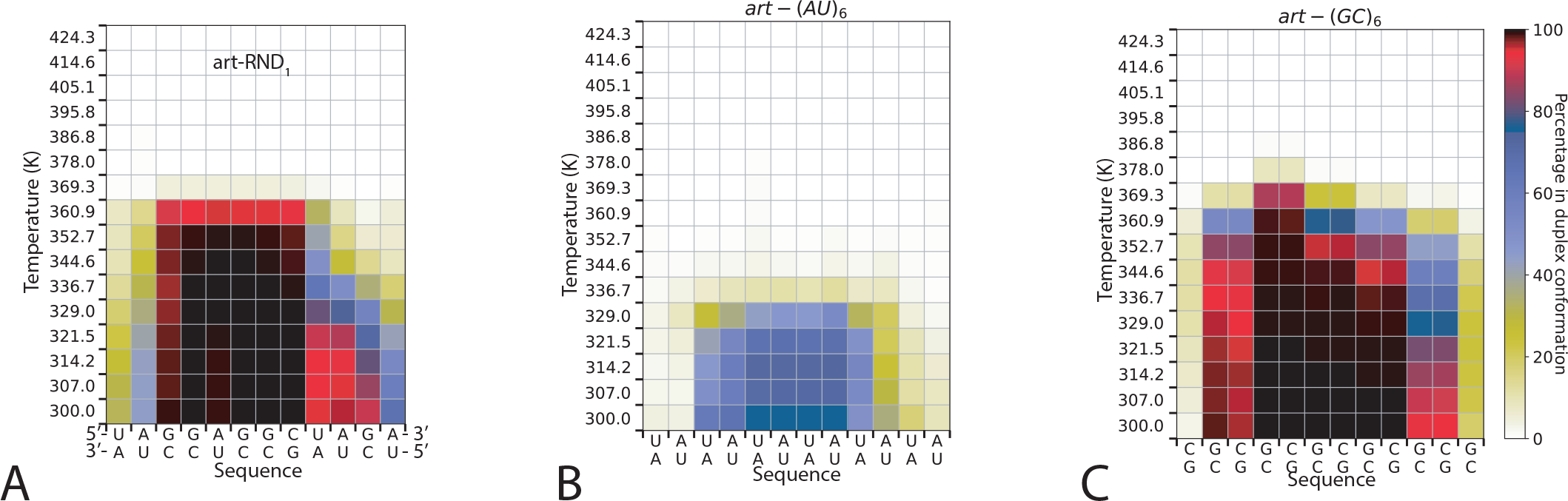
Local duplex melting. Fraction of formed base-pairs (color scale) between the two complementary strands of artificial strands not shown in the main text.

## Notes

### Competing Interest Statement

The authors have declared no competing interest.

## References

(1) Elbashir, S. M.; Harborth, J.; Lendeckel, W.; Yalcin, A.; Weber, K.; Tuschl, T. Duplexes of 21-nucleotide RNAs mediate RNA interference in cultured mammalian cells. Nature 2001, 411, 494–498.

(2) Fire, A. RNA-triggered gene silencing. Trends Gen. 1999, 15, 358–363.

(3) Setten, R. L.; Rossi, J. J.; ping Han, S. The current state and future directions of RNAi-based therapeutics. Nat. Rev. Drug Disc. 2019, 18, 421–446.

(4) Wilson, R. C.; Doudna, J. A. Molecular mechanisms of RNA interference. Annu. Rev. Biophys. 2013, 42, 217–239.

(5) Szostak, J. W. The eightfold path to nonenzymatic RNA replication. J. Sys. Chem. 2012, 3, 2.

(6) Borer, P. N.; Dengler, B.; Tinoco, I.; Uhlenbeck, O. C. Stability of ribonucleic acid double-stranded helices. J. Mol. Biol. 1974, 86, 843–853.

(7) Freier, S. M.; Kierzek, R.; Jaeger, J. A.; Sugimoto, N.; Caruthers, M. H.; Neilson, T.; Turner, D. H. Improved freeenergy parameters for predictions of RNA duplex stability. Proc. Natl. Acad. Sci. U.S.A. 1986, 83, 9373–9377.

(8) Mergny, J. L.; Lacroix, L. Analysis of Thermal Melting Curves. Oligonucleotides 2003, 13, 515–537.

(9) Narberhaus, F.; Waldminghaus, T.; Chowdhury, S. RNA thermometers. FEMS Microbiol. Rev. 2006, 30, 3–16.

(10) Rinnenthal, J.; Klinkert, B.; Narberhaus, F.; Schwalbe, H. Direct observation of the temperature-induced melting process of the Salmonella fourU RNA thermometer at base-pair resolution. Nucleic Acids Res. 2010, 38, 3834–3847.

(11) Kortmann, J.; Sczodrok, S.; Rinnenthal, J.; Schwalbe, H.; Narberhaus, F. Translation on demand by a simple RNA-based thermosensor. Nucleic Acids Res. 2011, 39, 2855–2868.

(12) Weber, G. G.; Kortmann, J.; Narberhaus, F.; Klose, K. E. RNA thermometer controls temperature-dependent virulence factor expression in Vibrio cholerae. Proc. Natl. Acad. Sci. U.S.A. 2014, 111, 14241–14246.

(13) Guo, P. The emerging field of RNA nanotechnology. Nature Nanotech. 2010, 5, 833–842.

(14) Jabbari, H.; Aminpour, M.; Montemagno, C. Computational Approaches to Nucleic Acid Origami. ACS Comb. Chem. 2015, 17, 535–547.

(15) Dimitrov, R. A.; Zuker, M. Prediction of hybridization and melting for doublestranded nucleic acids. Biophys. J. 2004, 87, 215–226.

(16) Hawley, S. A. Reversible pressure– temperature denaturation of chymotrypsinogen. Biochemistry 1971, 10, 2436–2442.

(17) Becktel, W. J.; Schellman, J. A. Protein stability curves. Biopolymers 1987, 26, 1859–1877.

(18) Stirnemann, G.; Sterpone, F. Recovering protein thermal stability using all-atom Hamiltonian replica-exchange simulations in explicit solvent. J. Chem. Theo. Comput. 2015, 11, 5573–5577.

(19) Stirnemann, G.; Sterpone, F. Mechanics of protein adaptation to high temperatures. J. Phys. Chem. Lett. 2017, 8, 5884–5890.

(20) Mathews, D. H. Using an RNA secondary structure partition function to determine confidence in base pairs predicted by free energy minimization. RNA 2004, 10, 1178–1190.

(21) Spasic, A.; Berger, K. D.; Chen, J. L.; Seetin, M. G.; Turner, D. H.; Mathews, D. H. Improving RNA nearest neighbor parameters for helices by going beyond the two-state model. Nucleic Acids Res. 2018, 46, 4883–4892.

(22) Patel, D. J.; Hilbers, C. W. Proton Nuclear Magnetic Resonance Investigations of Fraying in Double-Stranded dApTpGpCpApT in H2O Solution. Biochemistry 1975, 14, 2651–2666.

(23) Moe, J. G.; Russu, I. M.Proton exchange and base-pair opening kinetics in 5’-d(CGCGAATTCGCG)-3’ and related dodecamers. Nucleic Acids Res. 1990, 18, 821–827.

(24) Snoussi, K.; Leroy, J. L. Imino proton exchange and base-pair kinetics in RNA duplexes. Biochemistry 2001, 40, 8898–8904.

(25) Sponer, J.; Bussi, G.; Krepl, M.; Banas, P.; Bottaro, S.; Cunha, R. A.; Gil-Ley, A.; Pinamonti, G.; Poblete, S.; Jurečka, P. et al. RNA structural dynamics as captured by molecular simulations: A comprehensive overview. Chem. Rev. 2018, 118, 4177–4338.

(26) Pinamonti, G.; Paul, F.; Noe, F.; Rodriguez, A.; Bussi, G. The mechanism of RNA base fraying: Molecular dynamics simulations analyzed with core-set Markov state models. J. Chem. Phys. 2019, 150, 154123.

(27) Zgarbová, M.; Otyepka, M.; Šponer, J.; Lankaš, F.; Jurečka, P. Base pair fraying in molecular dynamics simulations of DNA and RNA. J. Chem. Theo. Comput. 2014, 10, 3177–3189.

(28) Colizzi, F.; Bussi, G. RNA unwinding from reweighted pulling simulations. J. Am. Chem. Soc. 2012, 134, 5173–5179.

(29) Xu, X.; Yu, T.; Chen, S. J. Understanding the kinetic mechanism of RNA single base pair formation. Proc. Natl. Acad. Sci. U.S.A. 2016, 113, 116–121.

(30) Weber, G. Mesoscopic model parametrization of hydrogen bonds and stacking interactions of RNA from melting temperatures. Nucleic Acids Res. 2013, 41, 1–7.

(31) Ferreira, I.; Amarante, T. D.; Weber, G. Salt dependent mesoscopic model for RNA at multiple strand concentrations. Biophys. Chem. 2021, 271, 106551.

(32) Wienken, C. J.; Baaske, P.; Duhr, S.; Braun, D. Thermophoretic melting curves quantify the conformation and stability of RNA and DNA. Nucleic Acids Res. 2011, 39, e52.

(33) Jin, L.; Shi, Y. Z.; Feng, C. J.; Tan, Y. L.; Tan, Z. J. Modeling Structure, Stability, and Flexibility of Double-Stranded RNAs in Salt Solutions. Biophys. J. 2018, 115, 1403–1416.

(34) Sanstead, P. J.; Stevenson, P.; Tokmakoff, A. Sequence-Dependent Mechanism of DNA Oligonucleotide Dehybridization Resolved through Infrared Spectroscopy. J. Am. Chem. Soc. 2016, 138, 11792–11801.

(35) Jones, M. S.; Ashwood, B.; Tokmakoff, A.; Ferguson, A. L. Determining Sequence-Dependent DNA Oligonucleotide Hybridization and Dehybridization Mechanisms Using Coarse-Grained Molecular Simulation, Markov State Models, and Infrared Spectroscopy. J. Am. Chem. Soc. 2021, 143, 17395–17411.

(36) Zhang, J. X.; Fang, J. Z.; Duan, W.; Wu, L. R.; Zhang, A. W.; Dalchau, N.; Yordanov, B.; Petersen, R.; Phillips, A.; Zhang, D. Y. Predicting DNA hybridization kinetics from sequence. Nature Chem. 2018, 10, 91–98.

(37) Li, P. T.; Vieregg, J.; Tinoco, I. How RNA unfolds and refolds. Annu. Rev. Biochem. 2008, 77, 77–100.

(38) Hinckley, D. M.; Lequieu, J. P.; De Pablo, J. J. Coarse-grained modeling of DNA oligomer hybridization: Length, sequence, and salt effects. J. Chem. Phys. 2014, 141, 035102.

(39) Ouldridge, T. E.; Šulc, P.; Romano, F.; Doye, J. P. K.; Louis, A. A. DNA hybridization kinetics: zippering, internal displacement and sequence dependence. Nucleic Acids Res. 2013, 41, 8886–8895.

(40) Xiao, S.; Sharpe, D. J.; Chakraborty, D.; Wales, D. J. Energy Landscapes and Hybridization Pathways for DNA Hexamer Duplexes. J. Phys. Chem. Lett. 2019, 10, 6771–6779.

(41) Maciejczyk, M.; Spasic, A.; Liwo, A.; Scheraga, H. A. DNA duplex formation with a coarse-grained model. J. Chem. Theo. Comput. 2014, 10, 5020–5035.

(42) Perez, A.; Orozco, M. Real-time atomistic description of DNA unfolding. Angew. Chem. Int. Ed. Engl. 2010, 49, 4805–4808.

(43) Wong, K.-Y.; Pettitt, B. M. The Pathway of Oligomeric DNA Melting Investigated by Molecular Dynamics Simulations. Biophys. J. 2008, 95, 5618–5626.

(44) Piana, S. Atomistic Simulation of the DNA Helix-Coil Transition. J. Phys. Chem. A 2007, 111, 12349–12354.

(45) Zerze, G. H.; Stillinger, F. H.; Debenedetti, P. G. Thermodynamics of DNA Hybridization from Atomistic Simulations. J. Phys. Chem. B 2021, 125, 771–779.

(46) Dabin, A.; Stirnemann, G. Toward a molecular mechanism of complementary RNA duplexes denaturation. J. Phys. Chem. B 2023, 127, 6015–6028.

(47) Katava, M.; Kalimeri, M.; Stirnemann, G.; Sterpone, F. Stability and Function at High Temperature. What Makes a Thermophilic GTPase Different from Its Mesophilic Homologue. J. Phys. Chem. B 2016, 120, 2721–2730.

(48) Katava, M.; Stirnemann, G.; Zanatta, M.; Capaccioli, S.; Pachetti, M.; Ngai, K. L.; Sterpone, F.; Paciaroni, A. Critical structural fluctuations of proteins upon thermal unfolding challenge the Lindemann criterion. Proc. Natl. Acad. Sci. U.S.A. 2017, 114, 9361–9366.

(49) Maffucci, I.; Laage, D.; Stirnemann, G.; Sterpone, F. Differences in thermal structural changes and melting between mesophilic and thermophilic dihydrofolate reductase enzymes. Phys. Chem. Chem. Phys. 2020, 22, 18361–18373.

(50) Maffucci, I.; Laage, D.; Sterpone, F.; Stirnemann, G. Thermal Adaptation of Enzymes: Impacts of Conformational Shifts on Catalytic Activation Energy and Optimum Temperature. Chem. Eur. J. 2020, 26, 10045–10056.

(51) Lu, X.-J. 3DNA: a software package for the analysis, rebuilding and visualization of three-dimensional nucleic acid structures. Nucleic Acids Res. 2003, 31, 5108–5121.

(52) Abraham, M. J.; Murtola, T.; Schulz, R.; Páll, S.; Smith, J. C.; Hess, B.; Lindah, E. GROMACS: High performance molecular simulations through multi-level parallelism from laptops to supercomputers. SoftwareX 2015, 1-2, 19–25.

(53) Chen, Z.; Znosko, B. M. Effect of sodium ions on RNA duplex stability. Biochemistry 2013, 52, 7477–85.

(54) Zgarbová, M.; Otyepka, M.; Šponer, J.; Mládek, A.; Banáš, P.; Cheatham, T. E.; Jurečka, P. Refinement of the Cornell et al. Nucleic acids force field based on reference quantum chemical calculations of glycosidic torsion profiles. J. Chem. Theo. Comput. 2011, 7, 2886–2902.

(55) Joung, I. S.; Cheatham, T. E. Determination of alkali and halide monovalent ion parameters for use in explicitly solvated biomolecular simulations. J. Phys. Chem. B 2008, 112, 9020–9041.

(56) Jorgensen, W. L.; Chandrasekhar, J.; Madura, J. D.; Impey, R. W.; Klein, M. L. Comparison of simple potential functions for simulating liquid water. J. Chem. Phys. 1983, 79, 926.

(57) Berendsen, H. J. C.; Grigera, J. R.; Straatsma, T. P. The missing term in effective pair potentials. J. Phys. Chem. 1987, 91, 6269–6271.

(58) Zgarbová, M.; Luque, F. J.; Sponer, J.; Cheatham, T. E., 3rd; Otyepka, M.; Jurečka, P. Toward Improved Description of DNA Backbone: Revisiting Epsilon and Zeta Torsion Force Field Parameters. J. Chem. Theo. Comput. 2013, 9, 2339–2354.

(59) Zgarbová, M.; Šponer, J.; Otyepka, M.; Cheatham, T. E.; Galindo-Murillo, R.; Jurečka, P. Refinement of the Sugar– Phosphate Backbone Torsion Beta for AMBER Force Fields Improves the Description of Z- and B-DNA. J. Chem. Theo. Comput. 2015, 11, 5723–5736.

(60) Fröhlking, T.; Mlýnský, V.; Janeček, M.; Kührová, P.; Krepl, M.; Banáš, P.; Šponer, J.; Bussi, G. Automatic Learning of Hydrogen-Bond Fixes in the AMBER RNA Force Field. J. Chem. Theo. Comput. 2022, 18, 4490–4502.

(61) Steinbrecher, T.; Latzer, J.; Case, D. A. Revised AMBER Parameters for Bioorganic Phosphates. J. Chem. Theo. Comput. 2012, 8, 4405–4412.

(62) Mlýnský, V.; Kührová, P.; Zgarbová, M.; Jurečka, P.; Walter, N. G.; Otyepka, M.; Šponer, J.; Banáš, P. Reactive Conformation of the Active Site in the Hairpin Ribozyme Achieved by Molecular Dynamics Simulations with ε/ζForce Field Reparametrizations. J. Phys. Chem. B 2015, 119, 4220–4229.

(63) Izadi, S.; Anandakrishnan, R.; Onufriev, A. V. Building Water Models: A Different Approach. J. Phys. Chem. Lett. 2014, 5, 3863–3871.

(64) Tribello, G. A.; Bonomi, M.; Branduardi, D.; Camilloni, C.; Bussi, G. PLUMED 2: New feathers for an old bird. Comput. Phys. Commun. 2014, 185, 604.

(65) Essmann, U.; Perea, L.; Berkowitz, M. L. A smooth particle mesh ewald potential. J. Chem. Phys. 1995, 103, 8577.

(66) Berendsen, H. J.; Postma, J. P.; Van Gunsteren, W. F.; Dinola, A.; Haak, J. R. Molecular dynamics with coupling to an external bath. J. Chem. Phys. 1984, 81, 3684.

(67) Nosé, S. A unified formulation of the constant temperature molecular dynamics methods. J. Chem. Phys. 1984, 81, 511.

(68) Parrinello, M.; Rahman, A. Polymorphic transitions in single crystals: A new molecular dynamics method. J. Appl. Phys. 1981, 52, 7182.

(69) Roder, K.; Stirnemann, G.; Wales, J.; Pasquali, S. Structural transitions in the RNA 7SK 5 hairpin and their effect on HEXIM binding. Nucleic Acids Res. 2019, 1–17.

(70) Röder, K.; Stirnemann, G.; Faccioli, P.; Pasquali, S. Computer-aided comprehensive explorations of RNA structural polymorphism through complementary simulation methods. QRB Discovery 2022, 3, e21.

(71) Mlýnský, V.; Janeček, M.; Kührová, P.; Fröhlking, T.; Otyepka, M.; Bussi, G.; Banáš, P.; Šponer, J. Toward Convergence in Folding Simulations of RNA Tetraloops: Comparison of Enhanced Sampling Techniques and Effects of Force Field Modifications. J. Chem. Theo. Comput. 2022, 18, 2642–2656.

(72) Wang, L.; Friesner, R. A.; Berne, B. J. Replica exchange with solute scaling: A more efficient version of replica exchange with solute tempering (REST2). J. Phys. Chem. B 2011, 115, 9431–9438.

(73) Ouldridge, T. E.; Louis, A. A.; Doye, J. P. Extracting bulk properties of self-assembling systems from small simulations. J. Phys.: Condens. Matter 2010, 22, 104102.

(74) Dock-Bregeon, A. C.; Chevrier, B.; Podjarny, A.; Johnson, J.; de Bear, J. S.; Gough, G. R.; Gilham, P. T.; Moras, D. Crystallographic structure of an RNA helix: [U(UA)6A]2. J. Mol. Biol. 1989, 209, 459–474.

(75) Sheng, J.; Li, L.; Engelhart, A. E.; Gan, J.; Wang, J.; Szostak, J. W. Structural insights into the effects of 2’-5’ linkages on the RNA duplex. Proc. Natl. Acad. Sci. U.S.A. 2014, 111, 3050–3055.

